# Gut regulatory T cells mediate immunological tolerance in *Salmonella*-infected superspreader hosts by suppressing cytotoxic activity of T cells

**DOI:** 10.1101/2021.07.16.452606

**Authors:** LM Massis, S Ruddle, SM Brewer, R Schade, R Narasimhan, JD Honeycutt, THM Pham, JA McKenna, SW Brubaker, P Chairatana, JA Klein, M Margalef-Català, MR Amieva, JG Vilches-Moure, DM Monack

## Abstract

Superspreader hosts carry out most pathogen transmission events and are often disease tolerant since they remain asymptomatic despite high pathogen burdens. Here we describe the superspreader immune state that allows for disease tolerance. In a model of *Salmonella* infection, superspreader mice develop colitis with robust CD4^+^ and CD8^+^ T-cell responses, however, they remain asymptomatic. We found that superspreaders have significantly more regulatory T cells (Tregs) in the distal gut compared to non-superspreader infected hosts. Surprisingly, the depletion of Tregs did not induce pathogen clearance but rather exacerbated weight loss, increased gut inflammation, and compromised epithelial intestinal barrier. This loss of tolerance correlated with dramatic increases in cytotoxic CD4^+^ and CD8^+^ T cells. Interestingly, CD4 neutralization in Tregs-depleted superspreaders was sufficient to rescue tolerance. Our results indicate that Tregs play a crucial role in maintaining immunologic tolerance in the guts of superspreader mice by suppressing cytotoxic CD4^+^ and CD8^+^ T-cell activities.

**AUTHOR SUMMARY:** Superspreader hosts are the main cause of disease transmission and a very important public health concern. Here, we evaluated the immunological tolerance of the *Salmonella* infected superspreaders in a mouse model. By manipulating Tregs, we demonstrated the immunological mechanism from the host to maintain health status and high pathogen burden. Tregs depletion in the superspreaders led to severe disease, with damage of the intestinal epithelia, and high morbidity without having any effect on shedding and systemic *Salmonella* burden. Furthermore, we demonstrated that the damage of the intestinal epithelia was related to cytotoxic activity of T cells. When Tregs were depleted, CD8^+^ T cells produced high levels of granzyme B and perforin. CD8^+^ T cells neutralization in Tregs depleted mice led to increased cytotoxic CD4^+^ T cells. Interestingly, neutralization of CD4^+^ T cells in the Tregs depleted mice led to a reduction in the CD8^+^ T cells producing granzyme B and it was sufficient to rescue host tolerance in this model. We demonstrate for the first time that cytotoxic CD4^+^ T cells damage the epithelial intestinal barrier and contribute to loss of tolerance in the context of a superspreader host. These findings open new perspectives to understand mechanisms of tolerance in the intestine of a superspreader host.

## INTRODUCTION

In contrast to resistant hosts, which control infection and minimize disease by reducing pathogen load, immune tolerant hosts can sustain higher pathogen loads without losing health (1). Importantly, immune tolerant hosts can become carriers and transmit disease. To exemplify the impact of immune tolerant hosts, the Pareto 80/20 rule has been applied to demonstrate that 20% of infected hosts can be responsible for 80% of diseases that may result in death (2). The 20% responsible for transmitting disease are called superspreaders due to their capacity to spread pathogens to other hosts and, for some diseases, remain asymptomatic. Thus, it is critical to better understand superspreaders in order to address this important public health threat and disease reservoir (3–5).

To understand the superspreader *Salmonella* host, previous work from our group demonstrated that 129X1/SvJ mice can be chronically infected with *Salmonella enterica* serovar Typhimurium (*S*Tm) (6) whereby 25-30% of the mice become superspreaders, shedding more than 10^8^ bacteria per gram of feces (7). Superspreader *S*Tm-infected mice are considered tolerant as they lack any signs of disease (e.g., ruffled fur, weight loss, or decrease in temperature) and shed enough bacteria to infect naive hosts (7, 8). Superspreader *S*Tm hosts also possess a unique systemic immune profile with a neutrophil-dependent blunting of systemic Th1 responses (9). However, the intestinal immunologic response involved in the maintenance of the tolerogenic state of superspreader hosts remains uncharacterized.

The host immune response plays a role in maintaining inflammation to a level that does not lead to detrimental damage to the host (6). Regulatory T cells (Tregs) play a role in protecting the host from severe inflammation by dampening CD4^+^ and CD8^+^ T cell responses during viral infections and establishment of commensalism (10–13). However, Tregs can also facilitate pathogen persistence (14). For example, Tregs have been shown to suppress T cell activation and it correlates with increases in bacterial burden during the early stages of a persistent systemic *S*Tm mouse infection, but not in later stages of the infection (15). Although the systemic immune response is very important for the host response to the infection, it is very relevant to study the intestinal immune response in the gut of superspreaders, as it is colonized by high levels of pathogen and the major source of *S*Tm transmission(7). Here, we show that Tregs in the infected mucosa are critical for maintaining the health of superspreader mice. We used a model of Tregs depletion in *S*Tm superspreader hosts and characterized the immunological cellular mechanism involved in tolerance. Our findings show that Tregs are required to suppress cytotoxic T cells in the colon. In the absence of Tregs, cytotoxic T cells cause loss of tolerance in the superspreader hosts by disrupting the intestinal epithelial barrier.

## RESULTS

### *Salmonella*-infected superspreader hosts have increased Tregs in the distal gut

To characterize the intestinal immune response to *S*Tm in the tolerant superspreader mice, we infected 129X1/SvJ mice orally with 10^8^ CFU of wild-type *S*. Typhimurium strain SL1344. Approximately 25-30% of the mice became superspreaders around 21 days post-infection (p.i.), shedding more than 10^8^ CFU of pathogen per gram of feces (Fig. 1a; Fig. S1a), similar to our previously published reports (7, 9, 16). Superspreader mice are tolerant to the high load of *S*Tm in the intestine as demonstrated by a lack of significant weight loss over the course of 28 days p.i. (Fig. S1b). To characterize immune cell populations in the guts of infected mice, colonic immune cells were extracted from non-superspreader mice (e.g., mice shedding 10^2^ - 10^7^ CFU/g feces) and superspreader mice (Fig. 1b and Fig S1a) 28 days p.i. and cell populations were analyzed by flow cytometry. As previously published by our group(7, 9), the superspreader mice had increased inflammation in the distal gut as shown by high percentages of neutrophils, dendritic cells, and macrophages (Fig. S1c) when compared to non-superspreaders and uninfected mice. Superspreader mice had increased CD4^+^ (Fig. 1c) and CD8^+^ T (Fig. 1d) cell percentages in the colon compared to non-superspreader mice. Th17 cells are important to impair the translocation of bacteria from the intestine to systemic sites (17), to characterize these cells in the gut of superspreaders, we measured RORγt (expressed in Th17 cells) CD4^+^ T cells. There was an increase in Th17 cells in superspreaders compared to non-superspreader mice (Fig. 1e). We then measured the systemic levels of *S*Tm in the superspreaders compared to non-superspreaders (Fig. S1d), and there was no difference in the systemic levels of *S*Tm, which reflects previous findings that Th17 cells impair bacterial translocation from the gut (17). Tregs are well described to suppress the activity of T cells (Reviewed in (18)), to address if Tregs play a role in the tolerance of superspreader STm infected hosts we measured the expression of the transcription factor Foxp3 (expressed in Tregs) in the CD4^+^ T cells. Superspreader mice had a significantly higher percentage of Tregs compared to non-superspreader mice (Fig. 1f). Indeed, the levels of Tregs in non-superspreader mice were not significantly different from uninfected control mice (Fig. 1f). Thus, Tregs levels positively correlated with CFUs in the feces of superspreader mice (Fig. 1g), and the correlation was even bigger when correlating Tregs with CFU from the mice that shed >10^7^ CFU/g of feces (Fig. 1h), suggesting that Tregs are important for maintaining tolerance in superspreader hosts.

**Figure 1.**
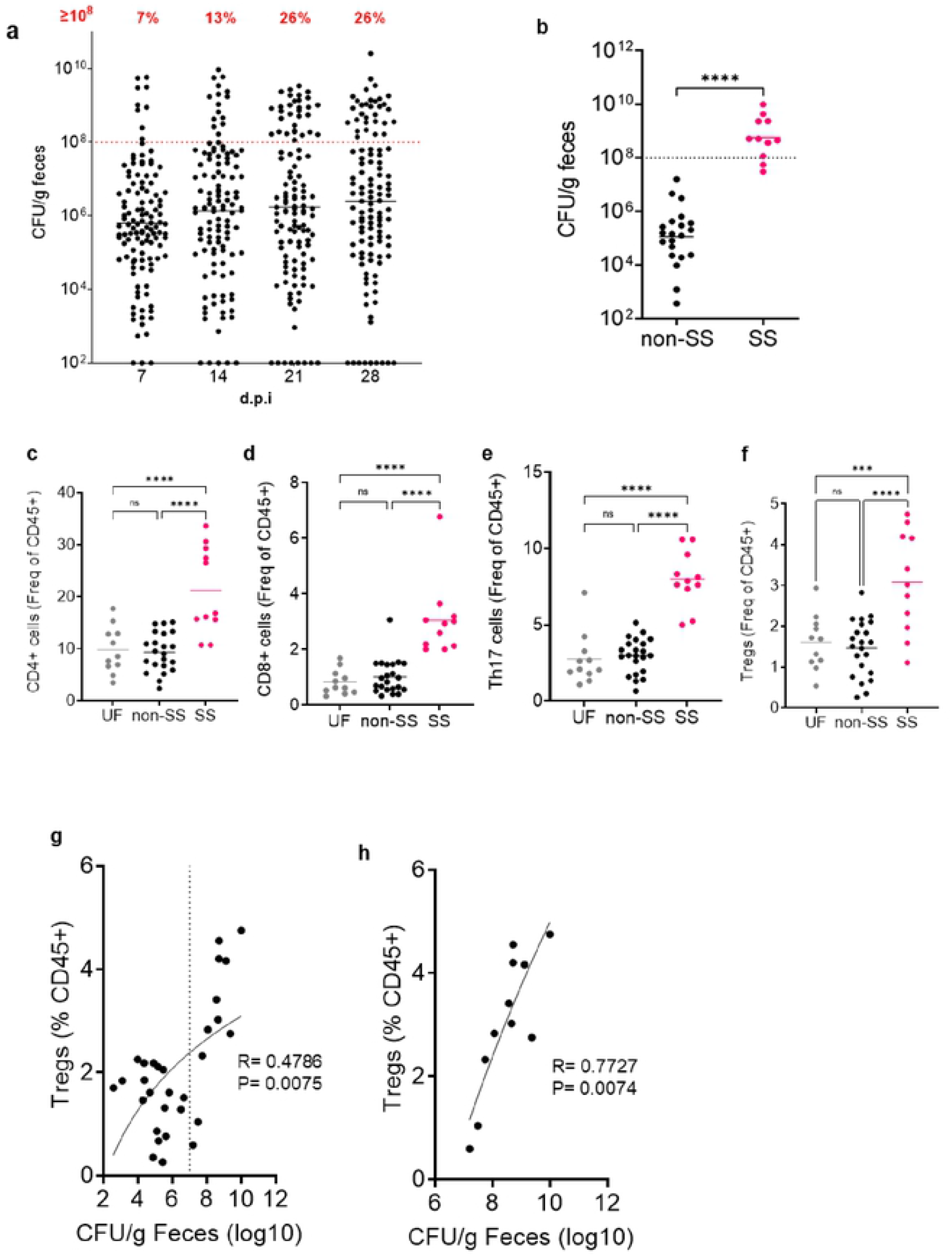
*Salmonella-infected* superspreader hosts have increased Tregs in the distal gut. 129×1/SvJ mice were infected with SL1344 by oral route (n=118, pooled from 2 independent experiments), **a**, CFU in the feces of infected mice over the period of 28 days. The red dashed line represents superspreader levels. The numbers in red are the percentage of SS mice by time point, **b**, Fecal CFU levels of Salmonella in the mice used for the colonic prep at day 28. Frequencies of CD45^+^ cells in the colon: CD4^+^ (**c**), CD8^+^ (**d**), CD4^+^Rorγt^+^ (**e**), CD4^+^Foxp3^+^ (**f**). **g**, Correlation of fecal spreading and Tregs in the colon, **h**, Correlation of fecal spreading and Tregs from the mice spreading more than 10^7^ CFU/g feces. Spearman’s rank correlation coefficients (R) and P two-tailed (P) are shown, (n= 21 non-SS, 11 SS, 10 UF). Data pooled from 2 independent experiments. Statistics: **b**, Mann-Whitney test, **c-f**, Data are presented as mean with each individual sample, and One Way ANOVA was used for comparison, **g-h**, Correlation line: Semilog line - X is log, Y is linear. non-SS-non-superspreader, SS-superspreader, UF-Uninfected controls. For all panels, P values less than 0.05 were considered significant (*P < 0.05; **P < 0.01; ***P < 0.001)

### Antibiotic-induced superspreader hosts phenocopy the natural superspreader immune response

Previous studies have shown an important role for Tregs in controlling the immune response to infections and maintaining homeostasis in the gut (19). In order to test the role of Tregs in maintaining tolerance, we used our previously published model to induce superspreaders in the mice infected with a *S*Tm strain that is streptomycin-resistant (7, 16). Briefly, we introduced gastrointestinal dysbiosis by administering a low dose of streptomycin (5 mg/mouse) 14 days p.i. As expected, (7, 16) two days after streptomycin administration, almost all mice shed >10^8^ CFU/g of feces (Fig. 2a) without a significant difference in *S*Tm levels in the spleen (Fig. 2b). The induced superspreader hosts did not exhibit significant weight loss or show signs of morbidity after antibiotic treatment (Fig 2c). The antibiotic-induced superspreader hosts had significant increases in the percentage and the total number of CD4^+^ T cells in the colon at day 21 p.i. (Fig. 2d), and by day 28 p.i. (Fig. 2h) the percentage of CD4^+^ T cells was similar to the levels in non-induced superspreaders (Fig. 1f). The percentage of CD8^+^ T cells was not altered at days 21 or 28 p.i. (Fig. 2e,i). Importantly, the mice had increased percentages of Th17 cells (Fig. 2f,j) and Tregs (Fig. 2g,k) at days 21 and 28 p.i. when compared to the mice that did not received the antibiotic treatment. These data indicate that these induced superspreader mice can be used as a model to study tolerance mechanisms in the colon of superspreader hosts.

**Figure 2.**
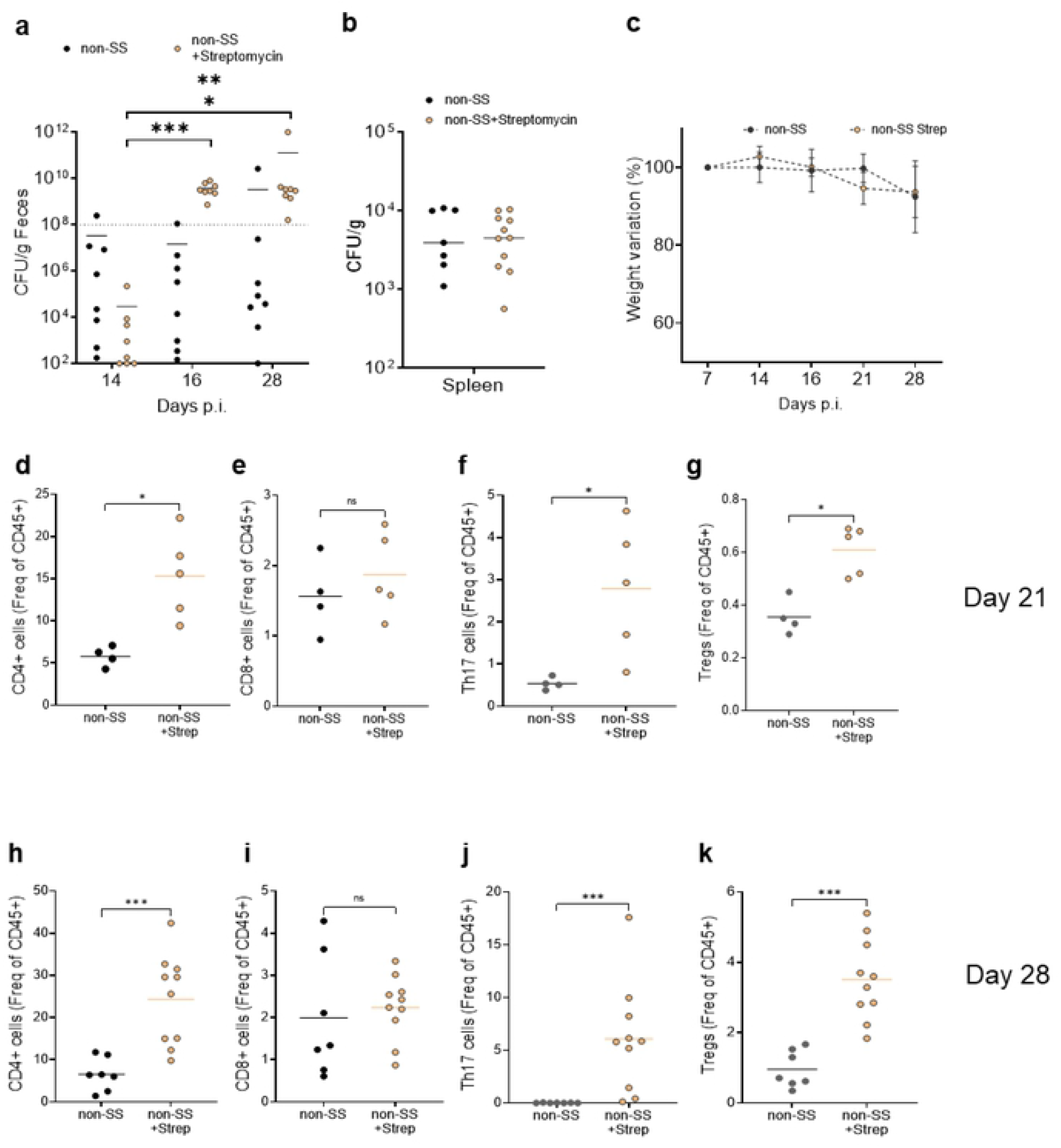
Antibiotic-induced superspreaders have increased Tregs in the colon. 129X1/SvJ mice were infected orally with 10^8^ SL1344 and treated with a low dose of Streptomycin at day 14 p. i. **a**, CFU/g of feces during the course of infection, **b**, CFU levels in Spleen at day 28 p.i. **c**, weight variation percentage starting at day 7 p.i. **d-k**, Colonic T cells assessed by flow cytometry. CD4^+^ cells in frequency of CD45^+^ cells in the colon at day 21 p.i. (**d**) and at day 28 p.i. (**h**). CD8^+^ cells in frequency of CD45^+^ cells in the colon at day 21 p.i. (**e**) and at day 28 p.i. (**i**). CD4^+^RORγt^+^ cells in frequency of CD45^+^ cells in the colon at day 21 p.i. (**f**) and at day 28 p.i. (**j**). CD4^+^Foxp3^+^ cells in frequency of CD45^+^ cells in the colon at day 21 p.i. (**g**) and at day 28 p.i. (**k**). n=9 mice-day 21 p.i., n=l8 mice-day 28 p.i. **a**, One-Way ANOVA, **d-k**, Mann-Whitney test. For all panels, P values less than 0.05 were considered significant (*P < 0.05; **P < 0.01; ***P < 0.001)

### Tregs depletion in superspreader hosts leads to loss of tolerance

To test the role of Tregs in the tolerance phenotype of superspreader hosts, we depleted Tregs in mice that express the high-affinity diphtheria toxin receptor under the control of the Foxp3 promoter which leads to efficient depletion of Tregs by diphtheria toxin (DT) injection (DEREG) (15, 20). Specifically, 129X1/SvJ males were crossed with FOXP3^DTR^ females to overcome Nramp1 susceptibility of C57BL6 hosts (21). Since the *Foxp3* gene is X-linked, the F1 129X1/SvJ X C57BL/6 male progeny have one copy of the *Foxp3*^DTR^ gene (F1-DEREG) and female progeny are *Foxp3* ^129X1/SvJ^ /*Foxp3* ^DTR^ (F1-DTR^+/−^) (Fig. 3a). The F1 progeny were infected with *S*Tm and the streptomycin treatment was used to induce the superspreaders. Tregs depletion was confirmed by the absence of Foxp3^+^ cells in the superspreader F1-DEREG mice (Fig. 3b,c) as well as the uninfected F1-DEREG mice (Fig. 3c). In addition, the percentage of CD4^+^ T cells (Fig. 3d) or Th17 cells (Fig. 3e) did not increase in Tregs-depleted mice. In contrast, CD8^+^ T cells increased in the colons of superspreader F1-DEREG mice compared to the levels of control superspreader F1-DTR^+/−^ mice (Fig. 3f), consistent with Tregs playing a role in suppressing CD8^+^ T cell levels (22). There was no change in the CD4^+^ and CD8^+^ T cell levels in uninfected F1-DEREG compared to uninfected F1-DTR^+/−^ (Fig. 3d,f).

**Figure 3.**
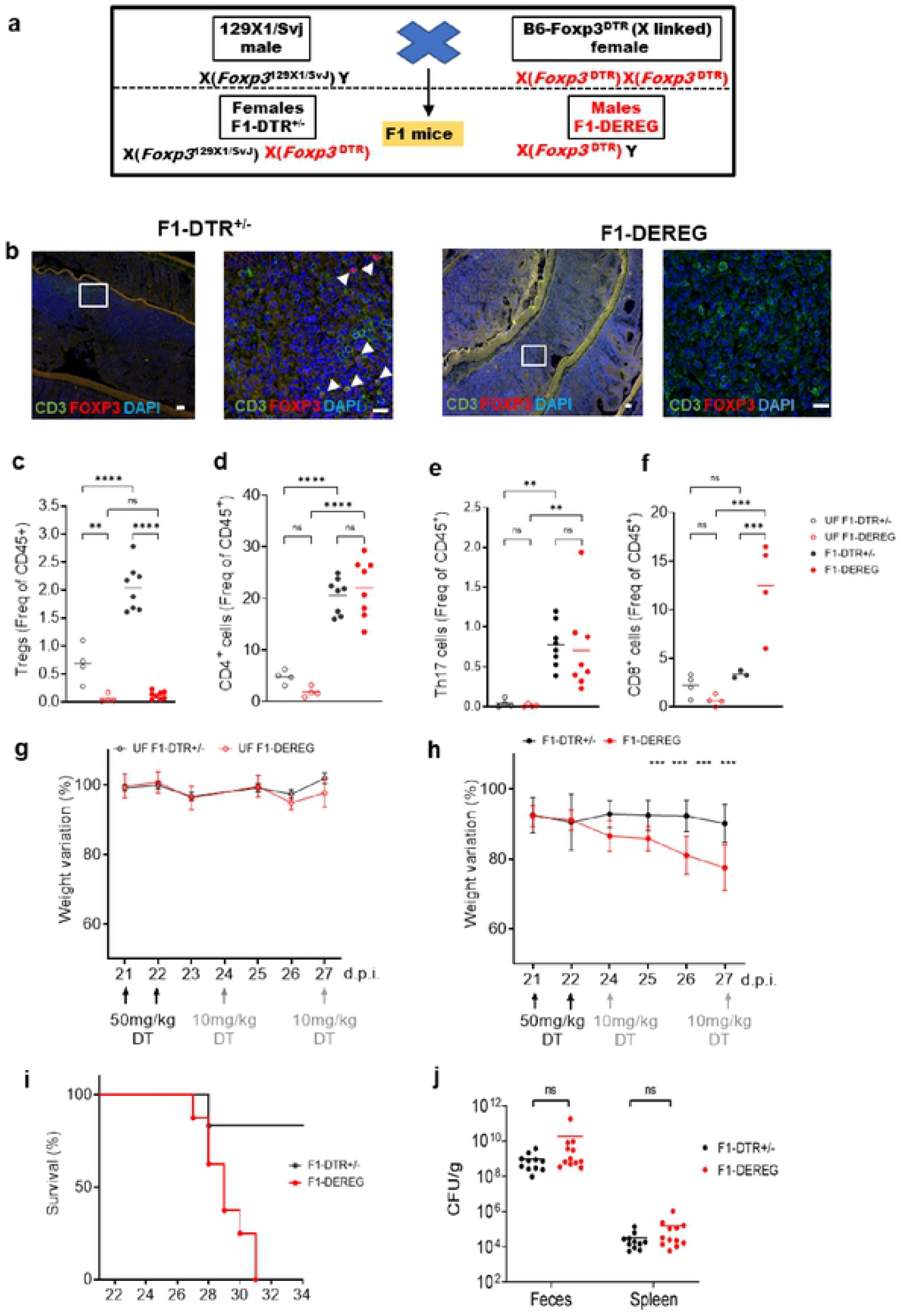
Depletion of Tregs leads to loss of tolerance and damaged intestinal epithelial barrier. **a,** Schematic representation of the F1-DEREG and F1-DTR^+/−^ mice breeding. F1-DEREG and Fl-DTR^+/−^ mice were orally infected with 10^8^ SL1344 or left uninfected, treated with a low dose of Streptomycin (5 mg) at day 14 p.i. and with diphtheria toxin (DT). **b**, Representative immunofluorescence staining of colonic “Swiss” roll with anti-CD3 Abs (green), anti-FOXP3 (red), and DAPI (blue). Arrowheads indicate Tregs. Scale bars are 50 μm (left panel) or 10 μm (right panel). Representative image of 3 mice in each group. CD4^+^ Foxp3^+^ (**c**), CD4^+^ (**d**), Th 17 (**e**), CD8^+^ (**f**) frequencies of CD45^+^ cells in the colon. Weight variation of F1-DEREG and F1-DTR+/− mice uninfected (**g**) or orally infected with 10^8^ SL1344 (**h**), treated with a low dose of Streptomycin (5mg) at day 14 p.i. and with diphtheria toxin (DT) as described in Methods, i, Survival curve of F1-DEREG and F1-DTR+/− mice infected with SL1344. **j**, CFU in the feces and spleen of infected mice. Mice numbers used: **b**, F1-DEREG n=25, F1-DTR^+/−^ n= 19, **c**, UF F1-DEREG n=12, UF F1-DTR^+/−^ n=l2, **d-f**, F1-DEREG n=8, F1-DTR^+/−^ n=8, UF F1-DEREG n=4, UF F1-DTR^+/−^ n=4, g, F1-DEREG n=12, F1-DTR^+/−^ n=11, **h**, F1-DEREG n=9, F1-DTR^+/−^ n=10, UF F1-DEREG n=11, UF F1-DTR^+/−^ n=11. **j**, F1-DEREG n=l2, F1-DTR^+/−^ n=11. Data pooled from 4 independent experiments, **i**, F1-DEREG n=9, F1-DTR^+/−^ n=7. Data from one experiment. Lines are mean. Statistic: **b-c,g**, Multiple Mann-Whitney test, **d-f,j**) One way ANOVA. UF, Uninfected. For all panels, P values less than 0.05 were considered significant (*P < 0.05; **P < 0.01; ***P < 0.001)

Finally, to address the role of Tregs in immunological tolerance in superspreader hosts, we investigated the influence of Tregs depletion on the severity of infection. Although all the control (Fig. 3g) and superspreader F1-DTR^+/−^ mice (Fig. 3h) (DT-treated uninfected F1-DEREG mice or *S*Tm-infected F1-DTR^+/−^ mice) showed only modest changes in body weight, Tregs-depleted superspreader (F1-DEREG) mice showed a significant drop in body weight within 3 days of DT treatment (Fig. 3h) and experienced increased morbidity, as indicated by ruffled fur, hunching, and malaise, by day 6 after Tregs depletion (Fig. 3i). To address the role of T cell responses in controlling pathogen levels in the absence of Tregs (15), *S*Tm burden was measured in the feces and spleen of F1-DTR^+/−^ and F1-DEREG superspreader mice. However, there was no change in the levels of *Salmonella* in the feces or the spleen of superspreader F1-DEREG mice compared to F1-DTR^+/−^ mice (Fig. 3j), indicating that the loss of tolerance in Tregs-depleted superspreader hosts is not due to increased pathogen burden. Rather, our results demonstrate that Tregs are important to maintain the immunological tolerance phenotype in superspreader hosts.

### Tregs depleted superspreader hosts have increased gut inflammation and loss of intestinal epithelial barrier function

To address possible mechanisms of Tregs-mediated tolerance, we characterized levels of inflammation and immune cells in F1-DEREG and F1-DTR^+/−^ *S*Tm superspreader mice. Similar to our previous studies (9, 16), the cecum (Fig. 4a, and S2a) and colon (Fig. 4b, and S2b) of superspreader mice were highly inflamed even in the presence of Tregs. The inflammation was characterized by infiltration of inflammatory cells within the intestinal wall, expansion of the connective tissue by edema fluid, and epithelial damage (Fig 4a,b,c). Importantly, there was a very significant increase in disruption of the epithelia, ulceration, and destruction of glands and crypts in the cecum and colon of superspreader F1-DEREG mice compared to superspreader F1-DTR^+/−^ mice (Fig. 4a,b,c). The cecum of uninfected F1-DEREG mice had higher pathology scores when compared to uninfected F1-DTR^+/−^ and the pathology cecum was generally greater than that in the colon (Fig.4c and Fig S2a), showing the importance of Tregs in the maintenance of intestinal integrity during homeostasis (Reviewed in (23)). To address the role of Tregs in the maintenance of immunological tolerance in the inflamed guts of superspreader mice, we measured the functional integrity of the intestinal epithelial barrier in the presence or absence of Tregs with a low-size (4kDa) FITC-Dextran assay as previously described (24, 25). Superspreader F1-DEREG mice had significantly higher levels of FITC-dextran in the plasma compared to the superspreader F1-DTR^+/−^ mice and uninfected control mice (Fig. 4d). Taken together, our data demonstrate that Tregs are crucial for maintaining intestinal epithelial integrity and maintaining tolerance in superspreader hosts.

**Figure 4.**
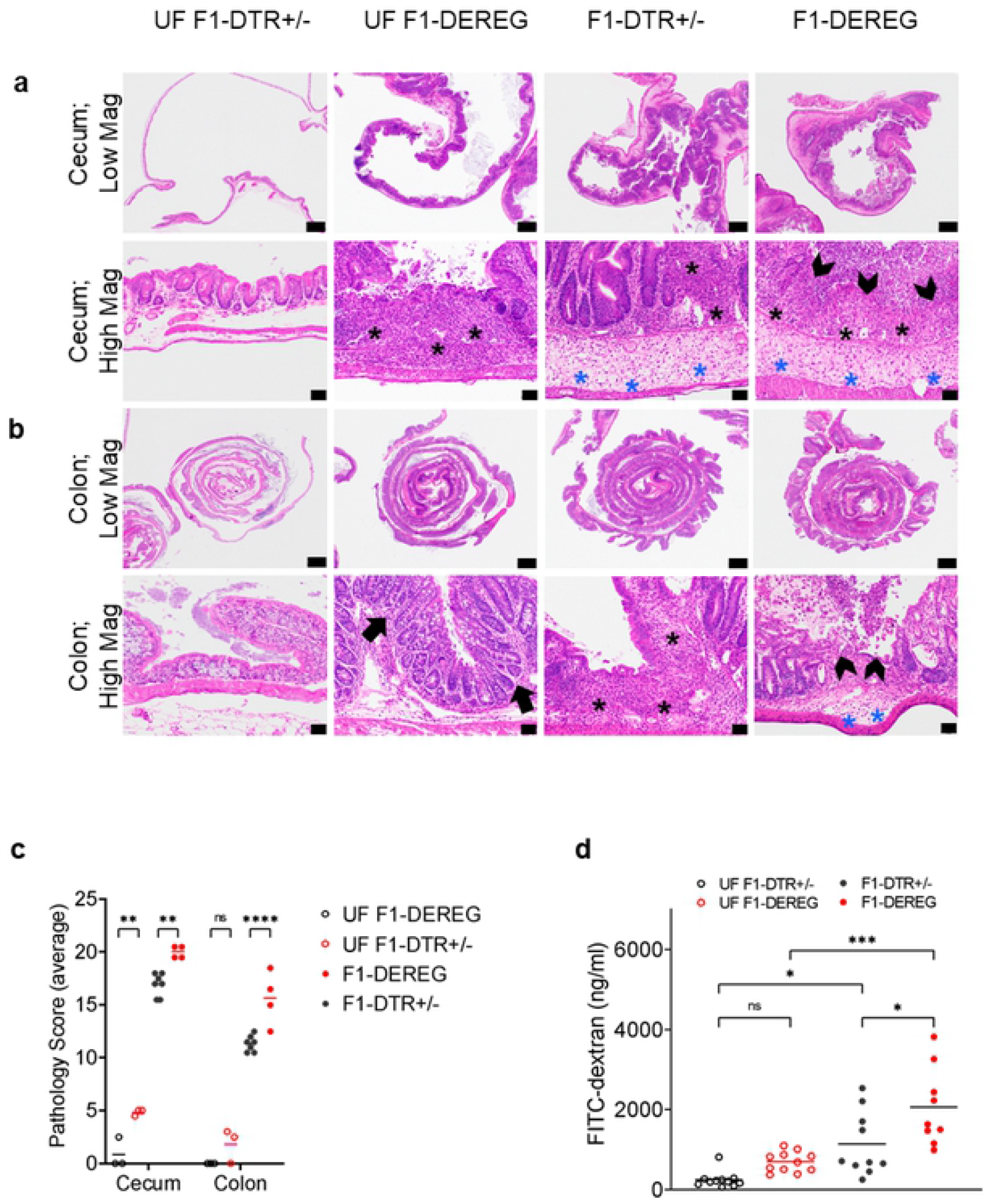
Depletion of Tregs increases the damage of the intestinal epithelial barrier. Low and high magnification images of hematoxylin and eosin (H&E)-stained sections of the cecum (**a**) and colonic “Swiss roll” (**b**) preparations. Upper panels= Magnification: l.25x; Scale Bars: 1.0 mm. Lower panels= Magnification: 2Ox; Scale Bars: 50 *μ*m. Representative image of 3 mice/group. **c**, Average of the pathology scores from colon and cecum, **d**, Gut permeability assay measured by FITC-dextran 4KDa in the plasma of the mice 28 days after infection. Statistics: Multiple Mann-Whitney tests. For all panels, P values less than 0.05 were considered significant (*P < 0.05; **P < 0.01; ***P < 0.001). Inflammatory cells in the lamina propria (black arrows). Paucity of glandular profiles and replacement by stroma and inflammatory cells (black asterisks). Submucosal edema (blue asterisks). Ulceration (black chevrons).

### Depletion of Tregs causes transcriptional changes in the colon of *S*Tm superspreader hosts

To gain more mechanistic insights into the role of Tregs in maintaining intestinal epithelial integrity in superspreader hosts, we performed gene expression analysis of the colons of infected mice. RNA was extracted from the proximal portion of the colon from F1-DEREG and F1-DTR^+/−^ mice infected for 28 days as well as uninfected mice. Previously published studies have shown that Tregs are important for epithelial wound healing (26, 27). To test the hypothesis that Tregs were functioning in wound healing pathways, we analyzed the transcriptional profile of the colons from F1-DTR^+/−^ and F1-DEREG mice. We used a NanoString nCounter gene expression system containing 770 genes that cover all the stages of tissue remodeling after injury (e.g., initiation of stress and immune responses cascades, inflammation, proliferation, and tissue remodeling) (Table S2 and Table S3). The transcriptional profiles in the colons of both *S*Tm-infected and uninfected mice in F1-DEREG mice were distinct from the F1-DTR^+/−^ as calculated by the log10 p-value and log2 fold change (Fig 5a). As expected, *Foxp3* expression is significantly higher in the F1-DTR^+/−^ infected or uninfected. However, there were no significant changes in the expression of genes involved with wound healing response, such as the receptor of Amphiregulin, EGF receptor (EGF-R) (28, 29) (Fig 5a, Fig S3f). In contrast, there was a significant difference in the expression of cytotoxic genes such as *Cd8a, Cd8b1, Prf1*, and *GzmB* in the colons of *S*Tm-infected superspreader F1-DEREG mice, suggesting that an increased cytotoxic response may be involved in damaging the gut epithelial barrier integrity (Fig. 5a, Fig S3b-e). To determine whether CD4^+^ and/or CD8^+^ T cells were involved in the cytotoxic response, the immune cells of the colon were extracted and the granzyme B (GZMB) levels were measured in CD4^+^ and CD8^+^ T cells by flow cytometry. The infected superspreader F1-DEREG mice had a higher percentage of CD8^+^ T cells (Fig. 5b) as well as a higher percentage of CD8^+^ T cells containing intracellular GZMB when compared to F1-DTR^+/−^ (Fig. 5c). In contrast, there was no change in the percentage of CD4^+^ T cells or of intracellular GZMB in the F1-DEREG infected mice when compared to F1-DTR^+/−^ infected mice (Fig. 5d,e). In addition, the presence of GZMB, perforin (PRF), and CD3^+^ cells in methacarn-fixed tissues were analyzed by immunofluorescence microscopy. Similar to our flow cytometry findings (Fig. 5c,e), colonic tissue sections from infected F1-DEREG mice contained significantly higher numbers of CD3^+^ cells co-staining with anti-GZMB and anti-PRF antibodies compared to colon sections from infected F1-DTR^+/−^ (Fig. 5f). The CD3^+^ cells of F1-DEREG mice stained with anti-GZMB and anti-PRF antibodies appear to be localized in clusters within the submucosal area of the colon (Fig. 5f), like previously described colonic patches (30). Collectively, our results suggest that Tregs play a role in dampening the cytotoxic CD8^+^ T cell-dependent epithelial damage in the tolerant superspreader hosts.

**Figure 5.**
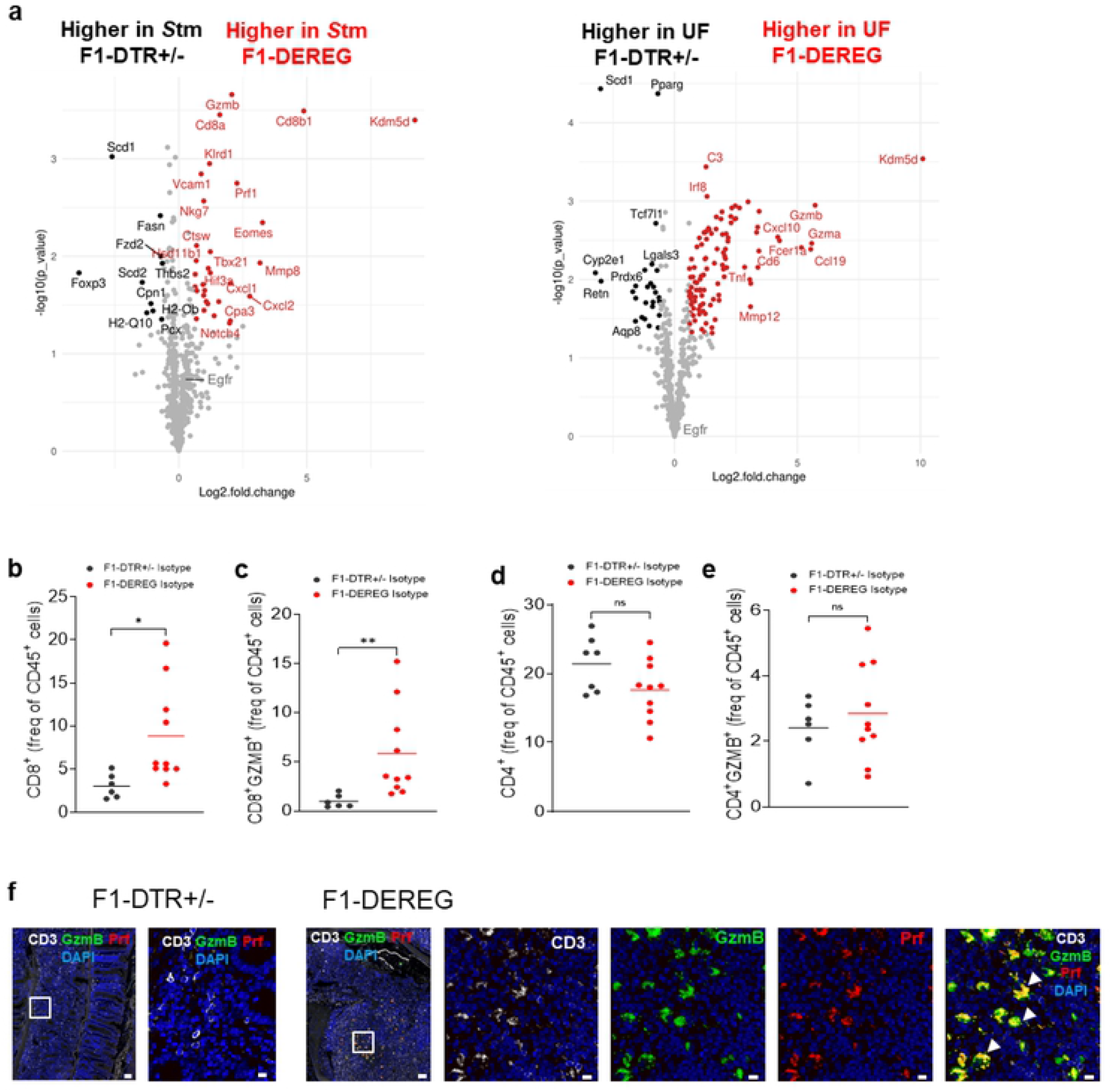
Tregs depletion alters cytotoxic response *during Salmonella* infection. **a**, Volcano plot of differentially expressed genes between colonic samples from F1-DTR+/− with F1-DEREG mice infected with SL1344 at day 28 p.i. (left panel) or uninfected (right panel), n= 3 per group. Mice were treated with DT as described in Methods. In red genes significantly highly expressed in F1-DEREG, in black genes significantly higher expressed in F1-DTR^+/−^, in gray genes not differentially expressed in F1-DTR^+/−^ or F1-DEREG. CD8^+^ (**b**), CD8^+^GzmB^+^ (**c**), CD4^+^ (**d**), CD4^+^GzmB^+^ (**e**) frequencies of CD45^+^ cells in the colon, **f**, Representative image of immunofluorescence staining of colonic “Swiss” roll with anti-CD3 Abs (white), anti-GzmB (green), anti-perforin (red), and DAPI (blue). Arrowheads highlight GzmB- and Prf-producing cells. Scale bars: 50 μm (left panel) or 10 μm (right panel). Representative image of 3 mice in each group. For all panels, P values less than 0.05 were considered significant (*P < 0.05; **P < 0.01; ***P< 0.001)

### CD4^+^ T cell neutralization restores tolerance in Tregs-depleted superspreader mice

To address the role of the cytotoxic T-lymphocyte (CTLs) in the loss of tolerance in Tregs-depleted superspreader hosts, we administered monoclonal anti-CD8b or isotype control antibodies to *S*Tm-infected superspreader F1-DEREG and F1-DTR^+/−^ mice as described in the Methods. We confirmed that CD8^+^ T cells were depleted in the colons of infected mice by flow cytometry (Fig 6a,b). Surprisingly, CD8^+^ T cell neutralization did not restore tolerance in F1-DEREG superspreader mice as weight loss was similar in the anti-CD8b cell antibody- and isotype control antibody-treated mice (Fig. 6c). In addition, the absence of CD8^+^ T cells did not alter the *S*Tm burden in the spleens and feces of superspreader F1-DEREG or F1-DTR^+/−^ mice (Fig. 6d). Finally, the level of gut epithelial damage was assessed with the FITC-dextran permeability assay in the CD8b-depleted mice and controls. The depletion of CD8^+^ T cells did not reduce gut permeability in the superspreader F1-DEREG mice (Fig. 6e), suggesting that other immune cell types contribute to loss of tolerance in these mice.

**Figure 6.**
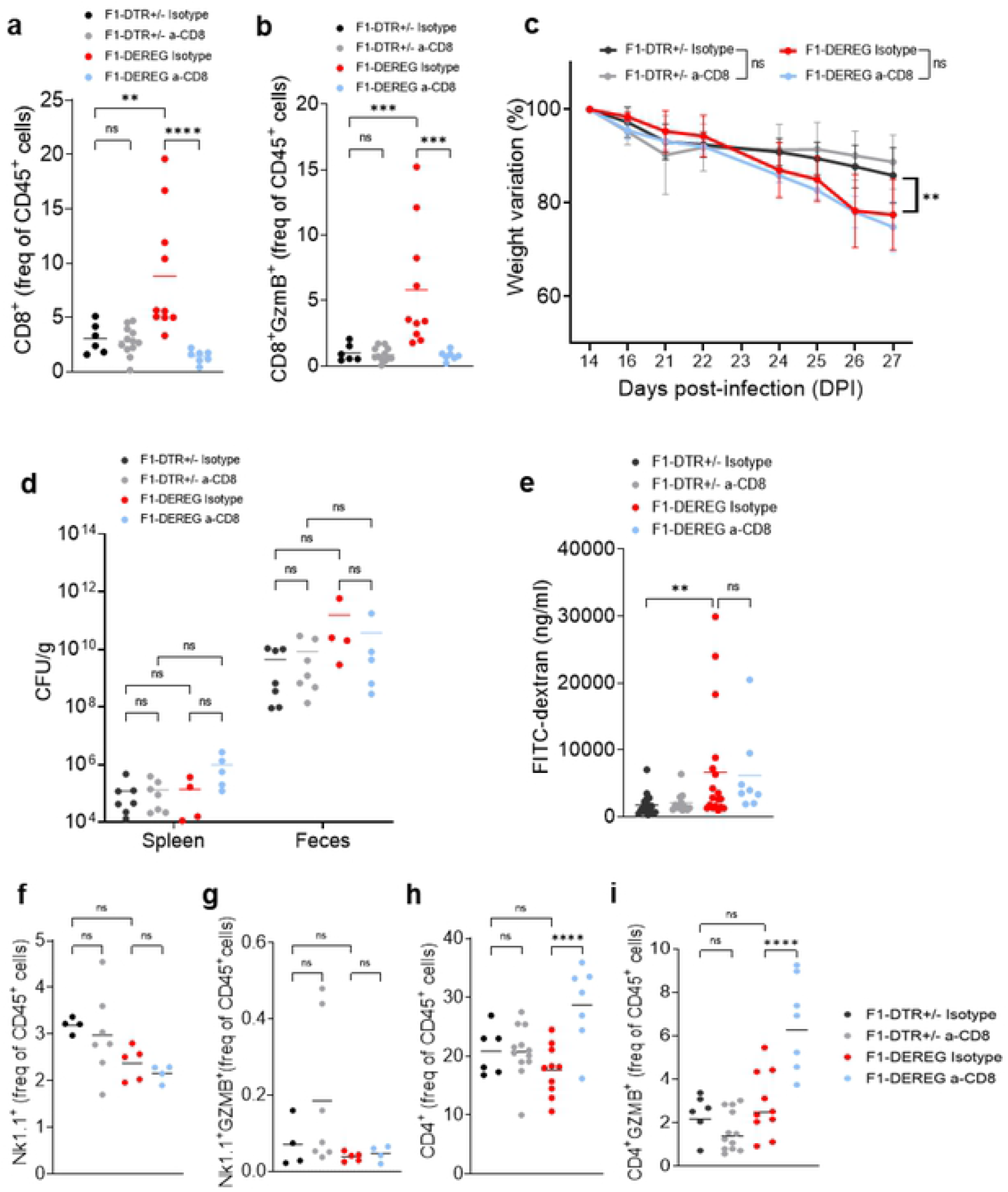
CD8^+^T cells neutralization induces CD4^+^T cells cytotoxicity in the colon of Tregs-depleted superspreaders. F1-DTR^+/−^ and F1-DEREG infected mice were treated with anti-CD8b (100 μg/mouse) or Isotype control (100 μg/mouse) as described in Methods. Colonic CD8^+^ (**a**), CD8^+^GzmB^+^ (**b**) T cells in percentage of CD45^+^ cells, **c**, Weight variation measured by the percentage of change from day 14 after infection, **d**, CFU levels per gram in spleen and feces, **e**, FITC-dextran levels in the plasma. Nk1.1^+^ (**f**), Nk1.1^+^ GzmB^+^ (g), CD4^+^ (**h**), CD4^+^GzmB^+^ (**i**) T cells in percentage of CD45^+^ cells in the colon. Statistics: Multiple Mann-Whitney tests. For all panels, P values less than 0.05 were considered significant (*P < 0.05; **P < 0.01; ***P < 0.001). N= 4-12 mice per group.

Therefore, we next investigated whether CD8^+^ T cell depletion would affect GZMB production in Natural Killer cells (NK) and CD4^+^ T cells. The percentage of NK cells in the CD8b-depleted mice did not increase (Fig. 6f), nor did the level of GZMB-producing NK cells increase when compared to isotype-control treated (Fig. 6g). However, there was a very significant increase in the percentage of CD4^+^ T cells in the colon of F1-DEREG mice treated with anti-CD8b when compared to the isotype treated (Fig. 6h). Interestingly, these CD4^+^ T cells acquired a cytotoxic phenotype as measured by the increased intracellular GZMB when CD8^+^ T cells were depleted (Fig. 6i). Collectively, our results indicate that cytotoxic CD4+ T cells are playing a major role in the loss of tolerance in Tregs-depleted superspreader hosts.

We next wanted to dissect the role of cytotoxic CD4+ T cells in loss of immune tolerance. Since neither an antibody to neutralize GZMB in vivo nor GZMB-deficient 129X1/SvJ mice are readily available, we neutralized both CD4^+^ and CD8^+^ T cells, or only CD4^+^ T cells in superspreader F1-DEREG and F1-DTR^+/−^ mice using neutralizing antibodies. The depletion was confirmed by flow cytometry (Fig 7e,g). Strikingly, F1-DEREG superspreader mice depleted of CD4^+^ and CD8^+^ T cells did not lose significant weight or show signs of morbidity, although mice that received the isotype control antibodies lost significant weight and became moribund (Fig. 7a). Surprisingly, *S*Tm-infected F1-DEREG mice depleted of only CD4^+^ T cells did not lose weight or show signs of morbidity (Fig. 7a).

**Figure 7.**
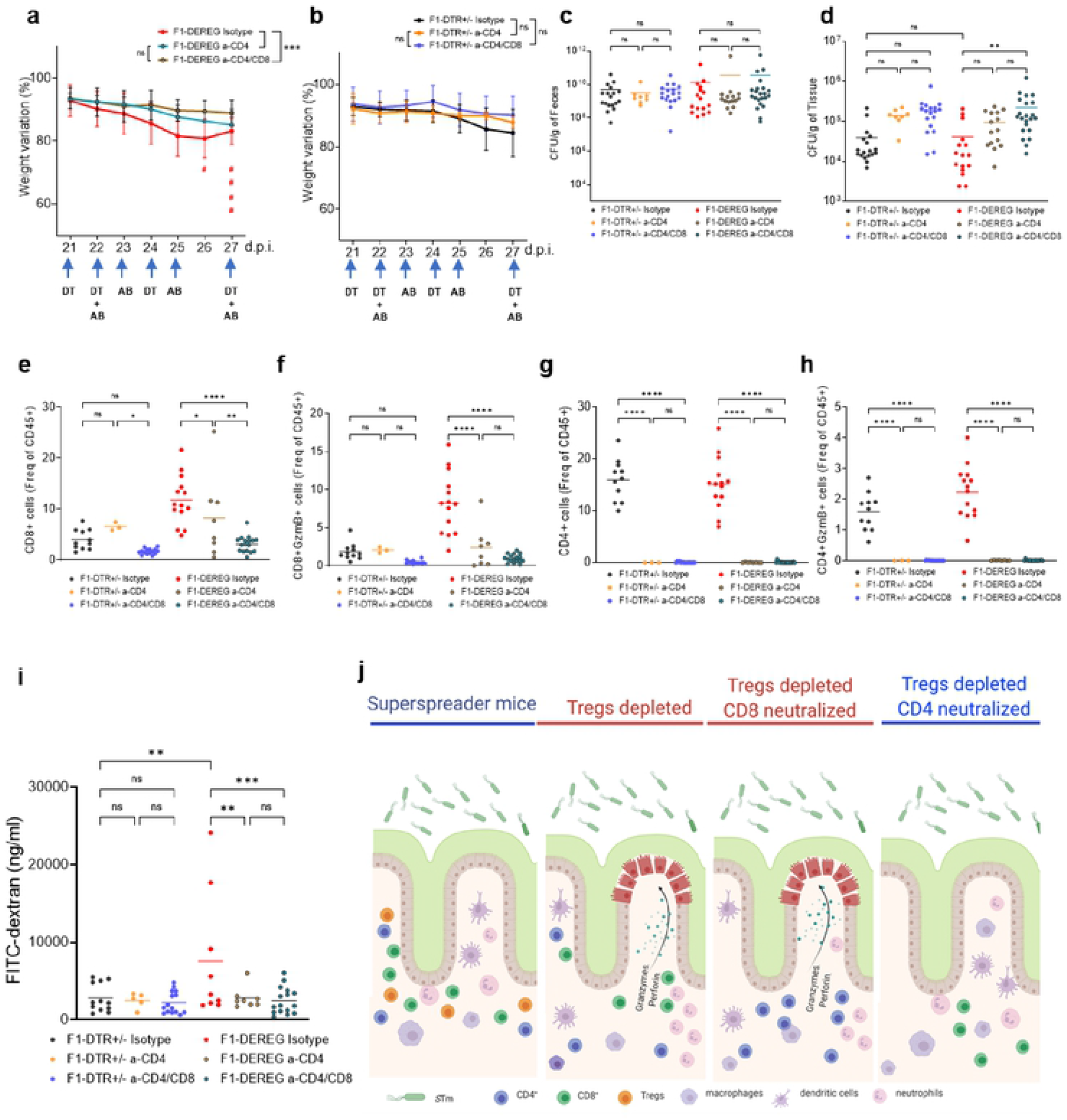
The tolerance of superspreader hosts is dependent on the cytotoxic response of T cells. F1-DTR+/− or F1-DEREG infected mice were treated i.p. with isotype-control (100 μg/mouse) and anti-CD4 (100 μg/mouse) or anti-CD4(100 μg/mouse) and anti-CD8(100 μg/mouse) or respective isotype control (100 μg/mouse of each isotype) as described in Methods, **a, b)** Weight variation measured by the percentage of change from day 14 after infection, red hash (#) represents each mouse that died from the Fl-DEREG Isotype group. CFU levels per gram in Feces **(c)** and Spleen **(d).** CD8^+^ **(e),** CD8^+^GzmB^+^ (f), CD4^+^ (g), CD4^+^GzmB^+^ **(h)** percentage of CD45^+^ cells in the colon measured by flow cytometry, **i**, FITC-dextran levels in the plasma, **j**. Schematic representation of the Tregs-dcpcndent supcrsprcader tolerance mechanism. Statistics: **a,b)** Multiple Mann-Whitney tests, **c-i)** One-way ANOVA. For all panels, P values less than 0.05 were considered significant (*P < 0.05; **P < 0.01; ***P < 0.001). a-d and f, N= 8-22 mice per group, g, Graphical abstracts created using Biorender.com.

Neutralization of just CD4^+^ T cells, as well as both CD4+ and CD8+ T cells in STm-infected F1-DTR^+/−^ superspreader mice, did not result in significant weight loss (Fig. 7b). Additionally, there was no change in the *S*Tm burden in the absence of CD4^+^ T cells or CD4^+^ and CD8^+^ T cells in the feces (Fig. 7c) and only slight increases in the spleen (Fig.7d) of superspreader F1-DEREG or F1-DTR^+/−^ mice. In addition, CD4^+^ T cell neutralization led to a decrease in the percentage of CD8^+^ T cells as well as intracellular GZMB levels in the colon of *S*Tm-infected F1-DEREG mice when compared to F1-DEREG mice (Fig. 7e,f). The neutralization of CD4^+^ T cell and GZMB intracellular levels was confirmed by flow cytometry (Fig. 7g,h).

To gain more insight into how CD4^+^ T cells might contribute to the greater weight loss of Tregs-depleted *S*Tm superspreader hosts, we analyzed transcriptional responses in infected F1-DEREG mice depleted of CD4^+^ and CD8^+^ T cells or CD4^+^ T cells utilizing the same NanoString nCounter gene expression system containing 770 that covers all the stages of tissue remodeling after injury. The relative expression of genes involved in the cytotoxic response was reduced in *S*Tm-infected F1-DEREG CD4^+^ and CD8^+^ T cell-depleted mice as well as *S*Tm-infected F1-DEREG CD4^+^ T cell-depleted mice. The normalized expression levels of *Cd8a, Cd8b, GzmB, Prf1* were significantly lower in the colon of CD4^+^, CD4^+^CD8^+^-neutralized *S*Tm-infected F1-DEREG mice compared to isotype control mice, confirming that the neutralizing antibodies dampened the cytotoxic response in the colon (Fig. S3a-f).

Importantly, the neutralization of both CD4^+^ and CD8^+^ T cells or only CD4^+^ T cells reduced gut permeability in the superspreader F1-DEREG mice to levels that were similar to superspreader F1-DTR^+/−^ mice (Fig. 7i), Taken together, our results indicate that Tregs-dependent suppression of CD4^+^ T cells in the guts of *S*Tm superspreader mice is crucial for immunological tolerance (Fig. 7j).

## DISCUSSION

Our work demonstrates that Tregs are a central player in controlling an exacerbated immune response that could be detrimental to the host. Understanding the mechanism of tolerance in a superspreader host is of great relevance since these hosts are responsible for most of the disease transmission. The present work shows how Tregs are a central player in controlling an exacerbated immune response that could be detrimental to the host.

Here, we demonstrate the role of Tregs in controlling cytotoxic T cell responses in the intestines of asymptomatic superspreader hosts. The depletion of Tregs increases the cytotoxic response induced by CD4^+^ T cells. These CD4^+^ T cells cause the activation of CD8^+^ T cells by releasing granzyme B and perforin. Our data suggest that the cytotoxic response causes greater damage to intestinal epithelia, disruption of the epithelial barrier, and loss of tolerance (Fig. 7g). To our knowledge, this is the first demonstration of this mechanism in the guts of a superspreader host.

In other infection models, the depletion of Tregs resulted in increased activation of T cells and control of the pathogen burden (15, 31–35). Johanns et al. (15) demonstrated that after intravenous infection with *S*Tm, Tregs play a tempo-dependent role in pathogen clearance. In our model, we orally infected mice, as this is the natural route for *Salmonella* infection. We checked the immune response and pathogen clearance both systemically (spleen) and in the feces of chronically infected and superspreader mice. Upon oral infection, Tregs depletion does not lead to pathogen control, but rather to gut permeability (as measured by FITC-dextran assay) and morbidity (as measured by weight loss).

Several other studies have shown that in the absence of Tregs, there is an increase in pathogen burden (36, 37). In the work by Wang et al. (36), Tregs depletion in a model of *Citrobacter rodentium* infection led to a higher pathogen burden and disease severity, but interestingly, lower histopathology score. In this study, we demonstrated that the depletion of Tregs in chronically infected superspreaders leads to an increase in inflammation, ulceration, and intestinal permeability, showing the protective role of Tregs in the context of a superspreader host.

Tregs depletion greatly impacts tolerance, as measured by weight loss and morbidity. To understand the mechanism of tolerance loss after Tregs depletion, we used a transcriptome-based approach to check for the expression of genes related to the steps of inflammation, healing, and tissue remodeling. The most prominent response found by this approach led us to investigate the cytotoxic response in the absence of Tregs. We confirmed the transcriptome data by flow cytometry and immunofluorescence, where we could find foci in the colon with CD3^+^ cells releasing granzyme B and perforin.

Surprisingly, neutralization of CD8^+^ T cells in *S*Tm superspreader mice depleted of Tregs was not enough to rescue the tolerogenic phenotype, which is the opposite of what has been described recently for viral infections (38). However, our results demonstrating increased cytotoxic CD4^+^ T cells in the guts of superspreader hosts in the absence of Tregs and CD8+ T cells are similar to previous studies showing that in the absence of CD8^+^ T cells, CD4^+^ T cells can acquire a cytotoxic phenotype (39) and anti-tumoral responses (40) (Reviewed in (41)). The crosstalk between T cells within the guts of superspreader mice is complicated but could involve sequestration of IL-2 by Tregs (40) and in the context of *S*Tm-infected superspreader hosts Tregs suppress cytotoxic CD4^+^ and CD8^+^ T cells. However, the increased levels of CD4^+^ T cells expressing granzyme B when Tregs and CD8^+^ T cells are depleted suggests that CD4^+^ T cells drive the cytotoxic response of CD8^+^ T cells. Consistent with this idea, we found that when CD4^+^ T cells are neutralized in Tregs-depleted superspreader mice, CD8^+^ T cells do not produce higher levels of granzyme B, and tolerance is rescued.

Altogether, these data demonstrate the importance of Tregs in dampening the cytotoxic response in the context of high pathogen loads in a superspreader host. Although the inflammatory response is fairly robust, Tregs play a crucial role in keeping cytotoxic T cell responses in check such that the gut epithelium does not get damaged and lose barrier function as a result of infection. To our knowledge, this is the first time Tregs have been shown to control the cytotoxic lymphoid response for the maintenance of tolerance within the intestine of a superspreader host.

## MATERIALS AND METHODS

### Mice

129X1/SvJ (Stock No: 000691) and B6.129(Cg)-Foxp3tm3(DTR/GFP)Ayr/J (also known as FOXP3^DTR^, Stock No: 016958) were purchased from The Jackson Laboratory. 129X1/SvJ were also bred in-house from mice purchased from Jackson Laboratory. F1 mice were generated by a cross of the female FOXP3^DTR^ with the male 129X1/SvJ (both purchased from Jackson Laboratory). The offspring females carry a copy of the *Foxp3* from the 129X1/SvJ and a copy of the Foxp3-DTR, the offspring males carry only a copy of the *Foxp3*-DTR (x-linked). All mice were housed under specific pathogen-free conditions. Water and food were provided *ad libitum*. Mice were acclimated for one week at the Stanford Animal Biohazard Research Facility prior to experimentation. All animal procedures were approved by Stanford University Administrative Panel on Laboratory Animal Care (APLAC) and overseen by the Institutional Animal Care and Use Committee (IACUC) under Protocol ID 12826.

### Infection, bacterial loads, and antibiotic dysbiosis

A single colony of *Salmonella* Typhimurium SL1344 (*S*Tm) was grown for 16 hours in LB supplemented with 200 mg/mL of streptomycin at 37°C and 200 x g agitation. The OD was adjusted to OD 1= 10^9^ CFU/mL. Bacterial pellets were washed twice with PBS and resuspended to a concentration of 5×10^9^ CFU/mL. 8-12 weeks mice were starved from food for 14 hours and infected by feeding the mice with 20 μL of the bacterial suspension (5×10^9^ CFU/mL) directly into the mouth using a pipette.

Infection was monitored by placing mice in individual boxes and collecting 1-2 fecal pellets in tubes containing 500 μL of PBS. The samples were weighed, serially diluted, and spot plated in duplicates on LB agar plates for CFU counting, and the counts were adjusted by the weight of the pellets. On day 28 p. i., the mice were humanely euthanized, and the organs were collected in PBS, dissociated with a pestle grinder, and serially diluted for CFU count. The counts were adjusted by the organ weight.

To induce dysbiosis, mice received 5 mg (25 μL of 200 mg/mL stock solution) of streptomycin 14 days after the *S*Tm infection, by drinking from a pipette tip.

### Tregs depletion and in vivo CD4, CD8 T cells neutralization

Mice were intraperitoneally (i.p.) injected with 50 mg/kg DT (diphtheria toxin, sigma cat. D0564) on days 21 and 22 after infection and with 10 mg/kg DT on days 24 and 27 after infection.

To neutralize T cells, mice were i.p. injected with anti-CD4 and/or anti-CD8 monoclonal antibodies. 100 μg of either/both InVivoMAb anti-mouse CD8β (Lyt3.2) (BE0223, BioXcell) and InVivoMAb anti-mouse CD4 (GK1.5) (BE0003-1, BioXcell) or isotype-matched control antibody (rat IgG2a isotype control, anti-trinitrophenol, BP0089; rat IgG2b isotype control, anti-keyhole limpet hemocyanin, BE0090, BioXcell) were diluted in PBS and administered on days 22, 23, 25, and 27 p. i.

### Intestinal permeability assay

Mice were administered fluorescein isothiocyanate (FITC)–dextran by oral gavage at 0.44 mg per gram of body weight. Four hours later, mice were humanely euthanized, blood was collected, and FITC–dextran concentrations in the plasma were measured via fluorescence spectrophotometry on Synergy HTX with an excitation of 485 nm (20 nm bandwidth) and an emission wavelength of 528 nm (20 nm bandwidth). Plasma from mice not administered with FITC-dextran was used to determine the background. FITC-dextran concentration was determined by a standard concentration curve (25).

### Colonic and splenic cell isolation and flow cytometry

The colon was harvested by cutting distal to the cecum and at the rectum. The fat and mesentery were removed, and the tissue was cut into 0.5cm pieces. Intestinal tissues were sequentially treated with HBSS containing 1 mM DTT at 37°C for 15 min and washed with 25 mM HEPES and 5% FBS at 37°C for 30 min, and then dissociated with gentleMACS dissociator (Mylteni) and dissociated in RPMI containing 0.167mg/mL liberase TL (Roche), 0.25mg/mL DNase I (Sigma), and 5% FBS with constant agitation at 37°C for 30 min. The cell suspension was further passed through a 100 μm cell strainer, followed by 70 μm and 40 μm cell strainers (Falcon). The cells were resuspended in FACS buffer (1X PBS with 2mm EDTA).

For flow cytometry, the cells were incubated for 5 min on ice with Fc Block (TruStain fcX anti-mouse CD16/32, Biolegend) and washed with 200 μL FACS buffer. Cells were stained on ice for 15 min in FACS buffer with a cocktail of Live/Dead Fixable Blue Viability Dye (ThermoFisher) and antibodies for surface antigens. The antibodies were purchased from eBiosciences, BD Bioscience, or BioLegend (Table S4).

For the staining of transcription factors, cells were resuspended in fixation and permeabilization buffer for 16 hours at 4°C (Foxp3 staining buffer set from eBioscience). Cells were washed in a permeabilization buffer and stained with transcription factor antibodies following the manufacturer’s protocol. For granzyme B staining, cells were stained for surface markers before fixation and permeabilization and then subjected to intracellular staining for 30 min according to the manufacturer’s protocol (Cytofix/Cytoperm buffer set from BD Biosciences). For instrument compensation, splenocytes isolated from naïve mice were stained with α-CD4 antibodies. Flow cytometry acquisition was performed on an LSR Symphony or LSRII.UV (BD Biosciences) and analyzed using FlowJo software (Tree Star) accordingly to gate strategy shown in Fig. S4.

### Microscopy

Tissues were harvested and fixed in Methacarn solution for 48 hours at room temperature and maintained in ethanol 70%. The cecum was fixed whole and bisected. The colon was linearized, and the lumen opened. The serosal surface of the colon was placed against a filter paper (Whatman, Cat. no. 100 090), and the linearized and flattened colon was submerged in Methacarn and allowed to fix. After fixation, the linearized colon was rolled into a “Swiss roll”. Tissues were routinely processed, embedded in paraffin, sectioned at 5.0 mm, and routinely stained with hematoxylin and eosin (H&E) or processed for immunofluorescence. Tissues stained with H&E were visualized with an Olympus BX43 upright brightfield microscope, and images captured using an Olympus DP27 camera and cellSens software. Tissues were assessed blindly by a board-certified veterinary pathologist. Criteria evaluated included ulceration (defined as the absence of epithelial lining exposing the lamina propria or deeper layers), inflammation (defined as the presence of inflammatory cells at various layers of the intestinal wall), and edema and/or fibrin exudation (defined as the expansion of the connective tissue with edema fluid and/or fibrin, with or without inflammatory cells). Based on these criteria, a grading scheme was developed using the following parameters (which were scored separately): 1) percent of surface area ulcerated, 2) severity of inflammation, 3) depth of inflammatory infiltrate, 4) percent area affected by inflammation, 5) absence of glands/crypts, and 6) percent area affected by edema and/or fibrin. For each tissue, each criteria was graded twice, and the scored averaged. The grading scheme and definitions of each score are shown in Table S1 (modified from (42)).

For immunofluorescence, the paraffin was removed by heating the slides at 70°C and washing twice with xylene at 70°C, 10 min. Followed by two washes with 100% ethanol, 5 min. The slides were washed twice with Antibody wash buffer (1x TBS IHC wash buffer (Millipore Sigma) with Tween 20 + 0.1% BSA) for 5 min at room temperature (RT), followed by 30 min incubation with blocking solution (1x TBS IHC wash buffer with Tween 20 + 2% Normal Donkey Serum, 0.1% Triton X-100, and 0.05% Sodium Azide). The slides were stained with anti-rabbit Foxp3, anti-rat perforin, anti-goat Granzyme B, anti-rat CD3 overnight at 4°C (Table S3). Slides were washed twice with wash buffer, then incubated for 2 hours at RT with the respective secondary antibody and DAPI. The slides were mounted with Prolong Diamond (Life Technologies) and cured for 48 hours before image acquisition. Images were collected using a 10X or 40X oil immersion objective on a Zeiss LSM 700 confocal microscope (Carl Zeiss) with ZEN 2.3 SP1 software (Carl Zeiss) and processed using Volocity Image Analysis software (Quorum Technologies).

### RNA extraction and NanoString^®^ nCounter assay

Total RNA was extracted from colonic tissues using the RNAeasy Kit (Qiagen) following the manufacturer’s instructions and quantified by NanodropA total of 25 ng of RNA was used for NanoString nCounter assay and the codeset for Mouse Fibrosis Panel (Nanostring) was utilized. The hybridization, processing, and acquisition were performed at the Nanostring facility (NanoString Technologies, Seattle, WA). The normalization and differential expression analysis were conducted using NSolver 4.0 software (Nanostring).

### Statistics

All statistical analyses were performed in R (v. 4.0.3.) and GraphPad Prism 9.0 (GraphPad Software, Inc., San Diego, CA), and visualized with ggplot2 and prism. Differences between groups were considered to be significant at a p-value of <0.05.

## ACKNOWLEDGEMENTS

The authors thank members of the Monack and Amieva laboratories for valuable discussion and technical support as well as Bokai Xhu for the protocol for immunofluorescence on methacarn-fixed tissue. Tissues for brightfield histology were prepared by the Animal Histology Service (AHS). The graphical abstract was created with BioRender.com.

